# TM-Vec 2s: Accelerated Protein Remote Homology Detection

**DOI:** 10.64898/2026.02.05.704073

**Authors:** Aryan Keluskar, Paarth Batra, Valentyn Bezshapkin, James T. Morton, Qiyun Zhu

## Abstract

Understanding protein function is an essential aspect of many biological applications. The exponential growth of protein sequence data has created a critical throughput bottleneck for structural homology detection: While billions of protein sequences have been identified from DNA sequencing data, the number of protein folds underlying biology is surprisingly limited, likely numbering tens of thousands. The “sequence-fold gap” limits the success of functional annotation methods that rely on sequence homology, especially for newly sequenced genomes. TM-Vec is a deep learning architecture that can predict TM-scores as a metric of structural similarity directly from sequence pairs, bypassing true structural alignment. However, the computational demands of its protein language model (PLM) embeddings create a significant bottleneck for large-scale database searches. In this work we present TM-Vec 2s, a highly efficient model created through distillation from a foundational teacher model. Benchmarks on the CATH and SCOPe domains for large-scale database queries showed that TM-Vec 2s is 185× faster than the original TM-Vec, while achieving higher TM-score prediction accuracy. It is even more efficient than the structure-informed search implemented in Foldseek, while showing strong competition in identifying remote homology between protein molecules. TM-Vec 2s significantly expands scalable, efficient exploration of protein structural homology space.

## Introduction

Understanding protein function is critical for biological applications, including drug discovery, therapeutic development, and precision medicine. The ability to accurately assess structural relationships between proteins without experimental 3D structures enables applications including remote homology detection [1], functional annotation transfer [2], and evolutionary analysis [3]. Remote homology detection, wherein similarity between proteins exhibiting negligible sequence similarity (<0.25 identity) is identified through conserved three-dimensional architectures, constitutes a fundamental challenge in computational structural biology. This challenge is particularly acute for microbial genomes from understudied environments, where traditional sequence-based methods fail to identify functional homologs, hindering our ability to discover novel enzymes and understand microbial community function [4]. This gap has acquired increased relevance following the expansion of predicted structure databases [5].

Protein language models (PLMs) have emerged as powerful tools for modeling protein behavior [6, 7, 8]. State-of-the-art PLMs leverage transformer architectures adapted from natural language processing, learning to generate high-dimensional contextualized representations of amino acids that capture both local and long-range sequence dependencies. These representations have enabled remarkable advances in protein structure prediction, functional annotation, and mutation impact assessment [9].

The template modeling score (TM-score) [10] is the standard metric for quantifying structural similarity, providing a length-normalized measure that correlates well with fold similarity [11, 9]. By learning to embed proteins in a space where cosine similarity correlates with TM-scores, TM-Vec [1] achieved state-of-the-art performance in remote homology detection. It demonstrated that deep learning approaches can predict structural similarity directly from sequences with remarkable accuracy, something that was previously unattainable using sequence-based alignment techniques. Despite these advances, current PLM-based methods face a fundamental computational bottleneck. This makes large-scale database searches on the scale of millions or billions computationally infeasible, preventing their integration into routine bioinformatic pipelines. Structure-based alternatives like Foldseek [12] achieve faster search through the 3D interaction (3Di) structural alphabet, but require precomputed or predicted 3D structures, restricting their applicability to sequences without an additional sequence-structure alignment step. The exponential growth of biological sequence databases has generated a critical need for rapid and scalable methods capable of analyzing billions of protein sequences efficiently.

We address this computational bottleneck through training a distilled model, TM-Vec 2s. For this task, we evaluated a variety of foundational PLMs and integrated the highly efficient Lobster-24M model for fine-tuning, then train an intermediate model, TM-Vec 2. The intermediate is then distilled into TM-Vec 2s, which learns to embed sequences into a latent space where cosine similarity between Vectors correlates with TM-score. We present accuracy and performance benchmarks for this model, which will allow for fast and efficient remote homology detection to better understand unknown protein functions.

### Structural similarity prediction

The “gold standard” structural alignment algorithms, such as TM-align [11] and DALI [13], are computationally expensive as they directly superimpose 3D coordinates. Historically, when the throughput of experimental methods limited the accumulation of structural data, this computational overhead was negligible. However, the release of AlphaFold [14] and language-model-based predictors has catalyzed a massive data surplus, populating repositories like AlphaFoldDB and ESM with hundreds of millions of predicted macromolecular structures. The increase in data availability by several orders of magnitude has rendered existing methods computationally infeasible for large-scale analysis and necessitated the development of novel approaches.

Advances in natural language processing have allowed for novel methods that utilize 3Di structural alphabet as representation for alignment. Foldseek [12] addresses efficiency through algorithmic optimizations and the 3Di structural alphabet, achieving remarkable speed improvements while maintaining sensitivity comparable to traditional structural aligners. While these methods are highly accurate, they require precomputed or predicted structures, restricting their scalability.

### Protein language models for structure

The emergence of large-scale PLMs has transformed structural biology. ESM [7], ProtBERT [15], and ProtT5 [9] demonstrated that self-supervised learning on protein sequences effectively captures evolutionary and, hence, structural information. To verify the accuracy of these protein language embeddings, we rely on scores such as RNS, which assesses the confidence of each model’s embeddings given verified vs. random protein sequences to embed.[16] On this metric, we can see both ProtT5 and ESM2 outperform other models, though ProtT5 performs better on proteins with disordered regions. TM-Vec leveraged this by fine-tuning ProtT5-XL for structural similarity prediction, achieving superior performance compared to sequence-based methods. However, the computational cost of these large models has limited their practical implementation. Recent advances have produced smaller, more efficient protein language models; Lobster [17] achieved competitive performance with orders of magnitude fewer parameters through optimized training procedures. Ankh [18] demonstrated that careful architectural choices can maintain representation quality while reducing computational requirements. Further work has explored various efficiency improvements, such as parameter sharing [19, 20] and parameter-efficient fine-tuning (PEFT) [21], but few studies have addressed the task of efficient structural similarity prediction.

### Knowledge distillation in biology

Knowledge distillation [22] has shown promise in compressing large neural networks while preserving performance. Existing distillation techniques such as DEGU show that distilled models perform at the level of model ensembles, and even better than standalone models, for tasks in genomic deep learning.[23] In the protein language space, DistilProtBERT [24] successfully reduced ProtBERT’s size by 50% while maintaining accuracy in protein classification tasks. Another study showed that multi-teacher distillation can reduce time complexity and preserve accuracy across multiple tasks, including binary and mutli-class classification and regression [25]. However, for TM-score prediction, the challenge is not maintaining fine-grained numerical precision, but rather efficiently capturing global structural topology. While a key goal is distinguishing between different folds (TM-score ≥ 0.5) [26], the continuous regression output provides differentiation across degrees of similarity within and between folds.

## Results

### Computational efficiency

We benchmarked the efficiency of encoding sequences by the TM-Vec models using the CATH S100 protein sequence dataset (Fig. 1A, B, Table S1). Substantial improvement of encoding speed was observed from TM-Vec to the intermediate model TM-Vec 2, then to the distilled student model TM-Vec 2s. While TM-Vec costs 31.13 minutes to encode 50,000 sequences, TM-Vec 2s only requires 9.66 seconds, marking a 193× speedup. We also included Foldseek, a widely adopted protein structure search tool [12] in the comparison. A precomputed database of protein structures was obtained from the RCSB PDB [27] and fed into Foldseek. TM-Vec 2s is 2.19× as fast as Foldseek, which costs 21.11 seconds to encode 50,000 structures. Running Foldseek with GPU was slower than without in our test. When Foldseek was used with the ProstT5 protein language model [28] to enable direct generation of structural representations from sequences, the runtime increases to 24.47 minutes, making TM-Vec 2s 152× faster than it.

**Figure 1.**
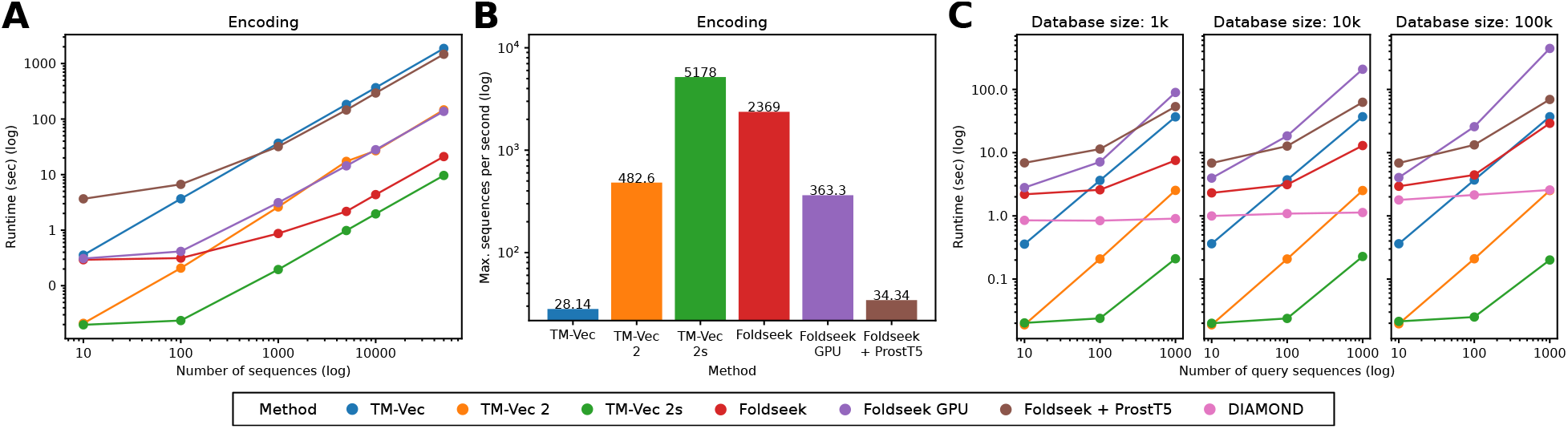
Runtime benchmarks for TM-Vec models compared with existing methods. **A**. Encoding time by the number of input protein sequences (TM-Vec models, Foldseek + ProstT5, DIAMOND) or structures (Foldseek, Foldseek GPU) (log-log scale). **B**. Maximum encoding speed per method from panel A, as represented by the number of sequences/structures encoded per second. **C**. Query time for 10, 100, and 1,000 query sequences/structures evaluated against databases of 1k, 10k, and 100k sequences.

We next benchmarked the efficiency of searching sequences against a pre-encoded database (Fig. 1C, Table S2). Since the TM-Vec models output embeddings, we used the FAISS libary [29] for similarity search of vectors. Significant acceleration was again observed in TM-Vec 2s versus the original TM-Vec. TM-Vec costs 37.04 seconds to search 1,000 sequences against a database of 100,000 sequences. In comparison, TM-Vec 2s only costs 0.20 seconds, which is 185× faster. Similar speedups were observed while querying against databases of various sizes. At the same time, TM-Vec 2s is 145× faster than Foldseek (29.12 seconds) or 346× faster than Foldseek with ProstT5 (69.44 seconds) in the above setting. It should be noted that in benchmarking Foldseek, we used the “search” option from its command-line interface, which performs a linear scan through the structural database without vector-based indexing, which potentially slows down the task. We also included DIAMOND, a widely used sequence similarity search tool [30] in the comparison. DIAMOND scales better over query size compared with the TM-Vec and Foldseek variants. Nevertheless, TM-Vec 2s is still 12.8× faster than DIAMOND in the above setting, or 85.4× faster when reducing query size to 100 sequences.

### Prediction accuracy

We evaluated all-vs-all TM-score prediction accuracy of TM-Vec models against reference TM-scores calculated by TM-align from structural alignments using the CATH S100 and SCOPe 40 datasets (Fig. 2, Table S3). From TM-Vec to TM-Vec 2s, notable increase of correlation strength (Pearson’s *r*: 8.4% in CATH S100 and 21.3% in SCOPe 40, respectively, same below; Spearman’s *ρ*: 12.4% and 21.8%) and reduction of prediction error (mean absolute error, MAE: 28.7% and 34.6%; root mean squared error, RMSE: 27.9% and 32.7%) were observed in both datasets (Fig. 2A-B, D-E), reflecting substantial improvement of TM-Vec 2s despite a much reduced model size. This is also supported by the distributions of TM-Vec 2s predictions being notably closer to ground truth compared with TM-Vec predictions (Fig. 2G).

**Figure 2.**
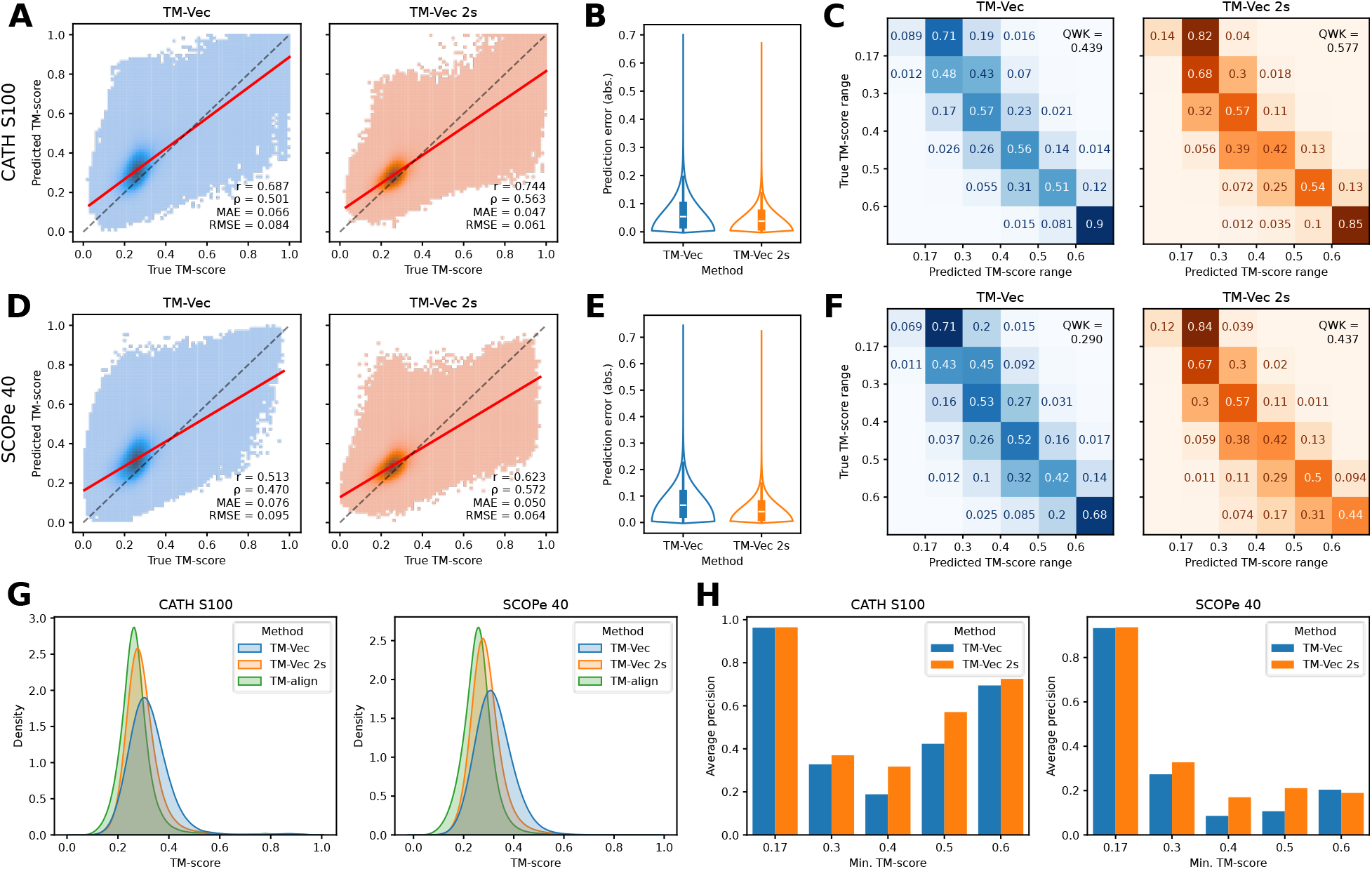
Comparison of TM-score prediction accuracy by TM-Vec models using CATH S100 and SCOPe 40 datasets. **A, D**. Correlation between the true TM-scores calculated by TM-align (*x*-axis) and predicted TM-scores by TM-Vec models, respectively (*y*-axis). Pearson correlation coefficient (*r*), Spearman correlation coefficient (*ρ*), mean absolute error (MAE) and root mean squared error (RMSE) are denoted. **B, E**. Distribution of absolute errors between predicted and true TM-scores. **G**. Distributions of TM-scores inferred by the three tools. **B, E** and **G**. The curves were computed using kernel density estimation (KDE). **C, F**. Confusion matrices showing the fractions of predictions for each of six TM-score ranges. Values less than 0.01 are not shown. Quadratic weighted kappa (QWK) is denoted. **H**. Average precision (AP) for binary classification divided by each of five TM-score bounds.

The regression plots reveal similar patterns of error distribution across the spectrum of TM-scores between the two models (Fig. 2A, D). The regression line crosses the diagonal line (true prediction) in the mid-range structural similarity (0.3 ≤ TM-score < 0.6). However, a closer look suggests that the TM-Vec 2s’ crossing point is closer to the lower side, while the deviation from truth is greater at the higher end, compared with TM-Vec, implying that TM-Vec 2s’ gain in low-to-mid-range prediction accuracy is accompanied with the reduced accuracy in high-range (TM-score ≥ 0.6, representing same structural families).

This observation motivated us to further diagnose the models’ strength and weakness by breaking down the [0, 1] range of TM-scores into six discrete bins. The two terminal bins, < 0.17 and ≥ 0.6, were defined to be consistent with the model design (they each received 5% training data; see above). The medium range, 0.17-0.6, which received 90% data during training, was further divided into four bins at the boundaries of 0.3, 0.4 and 0.5 [26], for higher granularity. The confusion matrices of per-bin classification reveal similar patterns again between the two models (Fig. 2C, F), with TM-Vec 2s being markedly better performing in assigning data to the correct bins (increase of quadratic weighted kappa, QWK: 31.5% for CATH S100, and 50.9% for SCOPe 40). An interesting observation is that neither model was competitive in assigning the lowest-similarity data (TM-score < 0.17, often representing random and unrelated protein pairs) to the correct bin. This pattern is consistent with the upticking left terminal of the regression line (Fig. 2A, D).

Additionally, we divided the TM-score spectrum into binary classes above or below each of the five boundaries defined above. The average precision (AP) scores calculated on these boundaries revealed notable improvement by TM-Vec 2s in the middle range (0.3, 0.4 and 0.5). Collectively, these observations reflect the improvement of TM-Vec 2s’ prediction accuracy overall, and particularly with protein domains with medium structural similarity, a “grey zone” that has been challenging to various categories of protein sequence and structure analysis methods.

### Homology detection

We evaluated homology detection performance across the CATH and SCOP structural hierarchies to assess model behavior as the task transitions from extremely remote (class) to near-homologous (family) relationships (Figs. 3, S4, Tables S4, S5). In addition to TM-align, we included the widely used protein structure search tool Foldseek [12] in the comparison. For comparability we utilize Foldseek’s exhaustive search threshold to maximize search result retention, and filtered all-vs-all protein domain pair TM-scores calculated by the other three methods to those reported by Foldseek. Homology detection is essentially a binary classification problem (positive if a pair of protein sequences belong to the same classification unit, negative otherwise), with highly imbalanced class sizes, with the positive class being much smaller but carrying more biological significance as they represent actionable research targets. We based our evaluation on the precision-recall (PR) curve and average precision (AP) (Figs. 3A, S4A), following the theoretical foundation [31, 32] (however see [33]) and common practices of the field. As expected, the structure aligner TM-align consistently outperforms the other three methods (with the exception of CATH class), reflecting the importance of accurate TM-score estimation for homology detection. The original TM-Vec consistently outperforms Foldseek. TM-Vec 2s’ performance surpasses TM-Vec in class and topology/fold levels in both datasets, while trailing closely in CATH architecture and superfamily levels. Meanwhile, it outperforms Foldseek in all six levels mentioned above. However, TM-Vec 2s’ performance notably decreased from TM-Vec at SCOP superfamily and family levels (AP dropped by 33.3% and 43.9%, respectively), and falls below Foldseek. This observation echos the reduction of high-similarity TM-score prediction accuracy described above.

**Figure 3.**
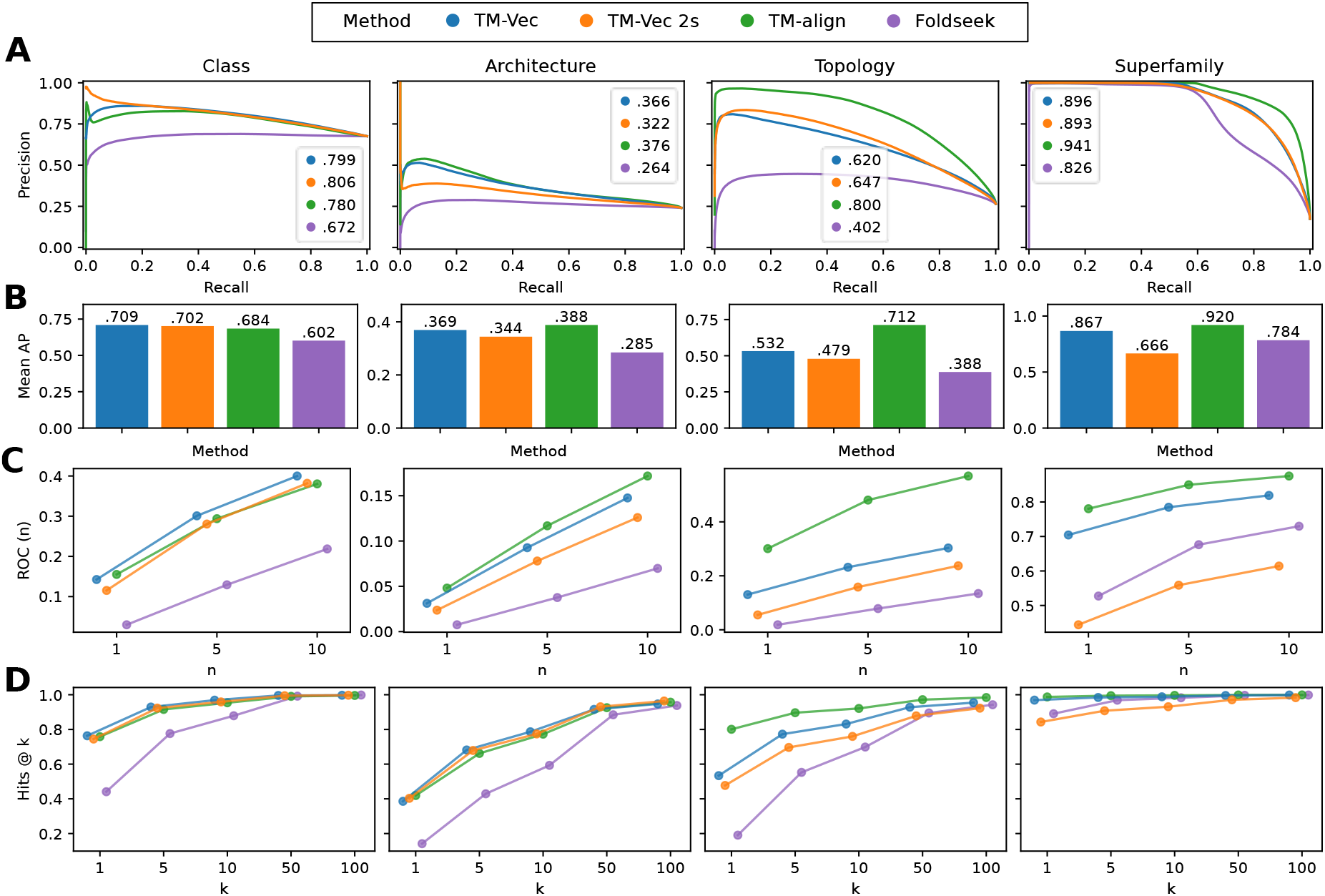
Comparison of homology detection capability of TM-Vec and TM-Vec 2s with reference to TM-align and Foldseek, using CATH S100 dataset at various classification levels (left to right: coarse to granular). Matches reported by all methods were retained. A match between a pair of protein domains is considered positive if they belong to the same classification unit, and negative otherwise. At each level, classification units with at least one positive and one negative were retained. Matches within the same lower-level classification unit were excluded. For example, topology/fold-level comparison exclude matches within the same superfamily. **A**.Matches were thresholded by the TM-score reported by each method to calculate the precision-recall (PR) curve and the average precision (AP, denoted in legend). **B**. Mean AP (MAP), i.e., macro average of APs of matches of each query. **C**. ROC_n_ (*k* = 1, 5, 10), i.e., matches were sorted by TM-score high-to-low within each query, and ROC_n_ is the ratio of true positives from the top of the match list down to the *n*-th false positive or the bottom, versus the number of actual positives of that query, averaged across queries. **D**. Hits@*k* (*k* = 1, 5, 10, 50, 100), i.e., fraction of queries within which the top *k* matches contain at least one true positive.

Structural classification units have highly uneven sizes, and larger units enriched in the training data may disproportionally influence AP values calculated from all protein domain pairs pooled together (i.e., micro average). To objectively assess model performance across a diverse spectrum of protein structures, we also calculated the mean AP over queries (MAP, aka macro AP) [34, 1, 35] (Figs. 3B, S4B). A similar but more consistent pattern was revealed, in which TM-align and TM-Vec consistently lead Foldseek, whereas TM-Vec 2s’ performance is close to TM-Vec and leads Foldseek at the topology/fold and above levels, but visibly decreased at the superfamily level and below.

Next, we compute the sensitivity up to the *n*-th false positive, ROC(*n*) [36], a metric frequently adopted to evaluate protein similarity search tools, as it prioritizes highly confident hits on top of the search results, which are most informative in a practical setting ([37, 38, 12]) (Figs. 3C, S4C). Similarly, TM-Vec 2s underperforms Foldseek at the superfamily level, but performs well at fold level. At the coarsest level, class, both TM-Vec models and TM-align demonstrate comparable performance, and significantly exceeds Foldseek. It’s also important to note that all measured tools underperform on ROC(n) when comparing structural similarity, as a range of factors impact the measurement of ground truth for this metric.

To more accurately compare the relevance of these metrics to measure remote homology, we calculate Hits@*k*, the proportion of queries in which the top *k* matches contain at least one true positive [39, 40] (Figs. 3D, S4D). This metric reflects practicality in biological research, where typically only a fixed number of candidates can be subjected to costly downstream analysis, and typically one success is all it needs to justify the study. All methods saturate at a large *k* number (100) at all levels, while the metric decreases substantially at smaller *k* numbers (10 and below). Likewise, TM-Vec 2s under-performs Foldseek at the superfamily and family levels, and outperforms at the topology/fold level and above. At the class and architecture levels, the two TM-Vec models and TM-align exhibit similar performance, and exhibit a substantial advantage over Foldseek.

To present a full image of this cross-method comparison, We also list the aforementioned performance metrics based on all protein domain pairs, without filtering to those reported by Foldseek (Figs. 3, S6, Tables S4, S7). As expected, metrics overall decreased, due to the inclusion of a large number of unrelated matches rather than enriching for detectable relationships. However, this analysis largely reproduced the patterns observed from the previous analysis and described above, showing TM-Vec 2s’ strong competence in predicting topology/fold-level and above relationships, at the cost of reduced performance with superfamily-level and below relationships.

## Discussion

We introduced TM-Vec 2s, a sequence-based model for protein structural similarity prediction that eliminates the computational barrier preventing deep learning methods from operating at database scale. By replacing the ProtT5-XL foundation model with the compact, efficient Lobster 24M model and distilling that knowledge into a lightweight BiLSTM architecture, we achieved speedups of up to 185× over the original TM-Vec for large-scale queries in our teacher model. Notably, TM-Vec 2s achieves the lowest mean absolute error across both benchmarks (MAE = 0.047 on CATH, 0.050 on SCOPe 40), indicating superior precision for fold discrimination despite slightly lower correlation, and out-performing the TM-Vec baseline.

The practical implication of these gains is a qualitative shift in structural bioinformatics. Unlike heuristic-based aligners like Foldseek, which only report scores for pairs passing sensitivity pre-filters, TM-Vec models provide continuous similarity predictions for every protein pair. This comprehensive search capability, combined with a training objective optimized for symmetric average TM-scores, allows our models to detect evolutionary signals in the “twilight zone” of structural similarity that are often invisible to coordinate-based or motif-based algorithms.

We hypothesize that since TM-Vec 2s was trained on a balanced dataset (with equal representation from the three TM-score bins) instead of 5/90/5% which TM-Vec 2 and TM-Vec were trained on, the student model learns decision boundaries in embedding space rather than over-fitting on detailed patterns, which may be more robust for fold classification.

It has been shown that there is large redundancy in the existing PLM latent space [17], and when we leverage distillation our models cause significant reduction in runtime without sacrificing accuracy when prediction structural homology from sequencing data.

Once encoded, TM-Vec embeddings enable amortized similarity search using vector indices like FAISS, allowing structural neighborhood exploration at the scale of billions of proteins. This positions TM-Vec 2s as a robust front-end for functional annotation and fold discovery, particularly in metagenomic settings where experimental structures are unavailable and predicted structures are computationally expensive to generate.

However, this speedup comes with trade-offs. Embedding methods provide only similarity scores, not residue-level alignments, which may limit their utility for downstream tasks requiring correspondence (e.g., close homology modeling, functional site transfer). Additionally, as learned approximations, they may fail on proteins with unusual structural features poorly represented in training data.[16]

While powerful, the current implementation has limitations. First, TM-Vec models cannot replace true structural alignment tools like TM-align where maximum accuracy is desired at the cost of lengthy runtime. Second, our benchmarks showed that TM-Vec 2s performs strongly in detecting remote homology whereas it underperforms alternative methods on closely related sequence pairs. We speculate that the distillation process retains higher-level resolution in the model, enabling it to detect remote homologies, while losing the resolution required to identify close relationships. The TM-Vec models also lack an associated statistical significance measure, such as *E*-values, which would enhance interpretability in massive database searches. Additionally, while the average TM-score provides stability, the impact of extreme query-target length asymmetry remains to be systematically characterized. Finally, extending these models from single-domain benchmarks to multidomain proteins will be essential to capture the full complexity of natural proteomes.

Despite these constraints, our results demonstrate that structural similarity can be inferred from sequences alone with higher sensitivity and greater efficiency than established structure-based methods. By removing the dependency on large protein language models and explicit structure generation, TM-Vec 2s enables structural reasoning to be applied as a routine operation. This shift bridges the long-standing gap between sequence abundance and structural insight, opening the door to structural annotation pipelines that scale with the modern genomic era.

## Methods

### TM-Vec 2: Identifying efficient foundation models

TM-Vec’s inference pipeline consists of three stages: 1) PLM embedding generation using ProtT5-XL, 2) siamese neural network for converting residual embeddings to structural embeddings, and 3) cosine similarity computation. We systematically evaluated three different foundation PLMs as alternatives to ProtT5-XL in the TM-Vec architecture: 1) Lobster [17] (24M and 150M parameters), an optimized transformer trained with highly efficient attention mechanisms; 2) Ankh [18], a larger 450M parameter model featuring architectural optimizations for robust protein sequence encoding; and 3) ProtMamba [41] (107M parameters), a state-space model that offers a distinct advantage over transformers by achieving linear time complexity with respect to sequence length.

We identified the foundation model which would be able to fine-tune on TM-Vec’s training dataset in the shortest duration (Fig. S1). The result showed that Lobster 24M is the fastest to fine-tune, completing in roughly 0.1 hours. Lobster 150M required more time, hovering around 0.15-0.2 hours, yet was still much quicker compared to the baseline ProtT5-XL model. ProtMamba, while promising in its architectural approach, lacked the hardware and community support to provide our desired speed, taking closer to 6 hours. We further benchmarked the inference times of the foundational models on various sequence length categories that represent the typical range of protein sequences encountered in structural biology applications (Fig. S2). Lobster 24M again exhibited the highest speed. Due to it’s strong performance for inference and in fine-tuning speed, we selected the Lobster 24M model for training TM-Vec 2 (Fig. 4).

**Figure 4.**
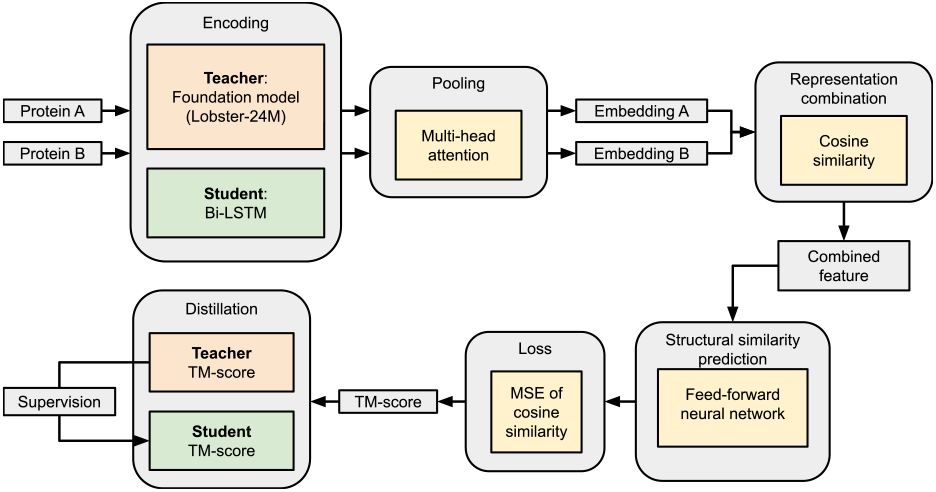
Illustration of the TM-Vec 2 teacher-student architecture. For each protein pair (*A, B*), the teacher model generates a TM-score. The student model encodes both sequences using a compact BiLSTM architecture and outputs a predicted TM-score. Training minimizes MSE loss that aligns the student’s predictions with the teacher’s outputs, enabling the student to approximate the structural topology learned by the teacher without using protein structures.

### TM-Vec 2s: Knowledge distillation architecture

While TM-Vec 2 reduces computational cost through foundation model replacement, the PLM embedding step remains a bottleneck. The TM-Vec 2s (TM-Vec 2 Student) model eliminates dependency on PLMs through direct knowledge distillation on raw amino acid sequences (Figs. 4, S3). Specifically, the target values are TM-scores computed by TM-Vec 2 between pairs of protein sequences derived from the ProteinNet dataset [42]. We use the *average* TM-score rather than the *maximum* TM-score for length normalization. This choice ensures symmetric scoring across protein pairs with significant length disparities, which is particularly important for remote homology detection where domain boundaries may differ. The student model contains 2.6 million parameters, a significant reduction of over 1,000× compared to ProtT5-XL, while maintaining comparable performance.

### Sequence encoder architecture

The student model employs a BiLSTM backbone [43], which has linear space complexity relative to sequence length, avoiding the quadratic complexity of self-attention in the transformer architecture [44, 45]. Each amino acid is mapped to a 128-dimensional embedding space, followed by a 2-layer bidirectional LSTM with 256 hidden units per direction (512 total). An attention pooling mechanism [46] aggregates the bidirectional hidden states into a fixed-length representation:

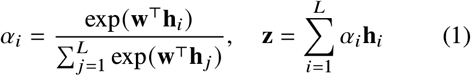

where **h**_*i*_ ∈ ℝ^512^ are the hidden states, **w** is the learned attention vector, and *L* is the sequence length. The pooled 512-dimensional vector passes through layer normalization to produce the final sequence embedding **z**. The output embedding **z** is kept 512-dimensional to maintain compatibility with the teacher model’s embedding space.

### TM-Score prediction

In the original TM-Vec architecture, for sequence embeddings **z**_*a*_ and **z**_*b*_, the TM-score is predicted via cosine similarity:

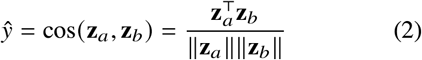

The student model maintains this characteristic by embedding proteins in a latent space where cosine similarity correlates directly with structural similarity. This enables the student model to support FAISS indexing [29] on the 512-dimensional embeddings for approximate nearest neighbor search.

### Training objective

The student model is trained to minimize the means-quared error in the embedding geometry:

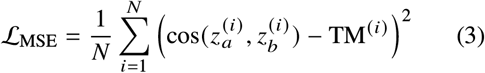

where *N* is the batch size, 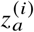 and 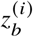 are the learned embeddings for protein pair *i*, and TM^(*i*)^ ∈ [0, 1] is the teacher model’s TM-score prediction. This direct regression objective stabilizes training and preserves the semantic structure of the teacher’s similarity space. The loss penalizes deviations between geometric distance in embedding space and structural distance in fold space.

### TM-Vec 2 training

We conducted training for the new TM-Vec 2 model by utilizing the DeltaAI cluster provided by National Center for Supercomputing Applications. Our dataset consists of 146 million pairwise protein comparisons derived from CATH v4.3.0 and aligned via TM-align. To prevent data leakage across folds, we employed Foldseek [12] (E-value < 0.01) to cluster available PDB structures. These clusters served as the basis for a train/test split, with 98% of clusters allocated for training and the remaining 2% reserved for testing. To ensure high representation of structurally similar proteins, the 146M pair dataset was adjusted to include a 5/90/5 split across three TM-score bins: 0-0.17 (low similarity, where there are predominantly random protein structural pairs), 0.17-0.6 (medium similarity, spanning from weak, above-random resemblance to probable fold-level similarity), and 0.6+ (high similarity, where most protein pairs belong to the same structural fold) [11, 26]. The model was trained on one NVIDIA GH200 node for 36 hours, and saw 10.6M samples per epoch, balanced according to the bin-based splitting strategy.

### TM-Vec 2s training

Training pairs for TM-Vec 2s were sampled from a data augmented pool of protein pairs used for TM-Vec 2 [42]. The pairs were first partitioned into the same three TM-score bins: low (0–0.17), medium (0.17–0.6) and high (0.6–1.0) as defined by TM-align [11]. An equal number of pairs was drawn from each bin. Within each bin, pairs were further divided into sub-intervals of 0.05, from which equal numbers of pairs were also drawn. The resulting three million pairs are therefore approximately uniform over the entire [0, 1] TM-score range. The target TM-score for training the student model was obtained using the TM-Vec 2 model. We have used a training/validation/test split of 64%, 16% and 20%, respectively. The student model was trained for 25 epochs on a system with one NVIDIA A100 GPU (80GB VRAM) for approximately 7 hours (with a batch size of 64 and AdamW optimizer using learning rate 1e-3, weight decay 0.01). We used Arizona State University’s Sol supercomputer [47] to conduct training for the TM-Vec 2s model.

### Hit retrieval verification

We ask whether TM-Vec 2 and TM-Vec 2s can guarantee a chosen error rate for incorrect hits. For each query we take its top *k* = 1000 neighbors, ranked by cosine similarity on *ℓ*_2_-normalized embeddings, and count a neighbor as a true hit if it has TM-Score ≥ 0.6.[4] The false-discovery rate (FDR) is the average fraction of returned hits that are wrong.

For a target FDR *α*, we pick a similarity threshold 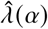 and keep only neighbors scoring above it. We choose the threshold with Learn-then-Test risk control (100 trials, *n*_calib_ = 1000, *δ* = 0.5, on a family-disjoint Pfam benchmark), which gives a finite-sample, distribution-free guarantee:

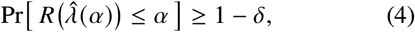

where *R* is the average per-query FDR. The empirical per-query value measured on held-out queries tracks the target *α* at every level (Fig. 5A): for example 0.095 ± 0.003 at *α* = 0.10, and similarly at *α* = 0.001, 0.01, and .50.

**Figure 5.**
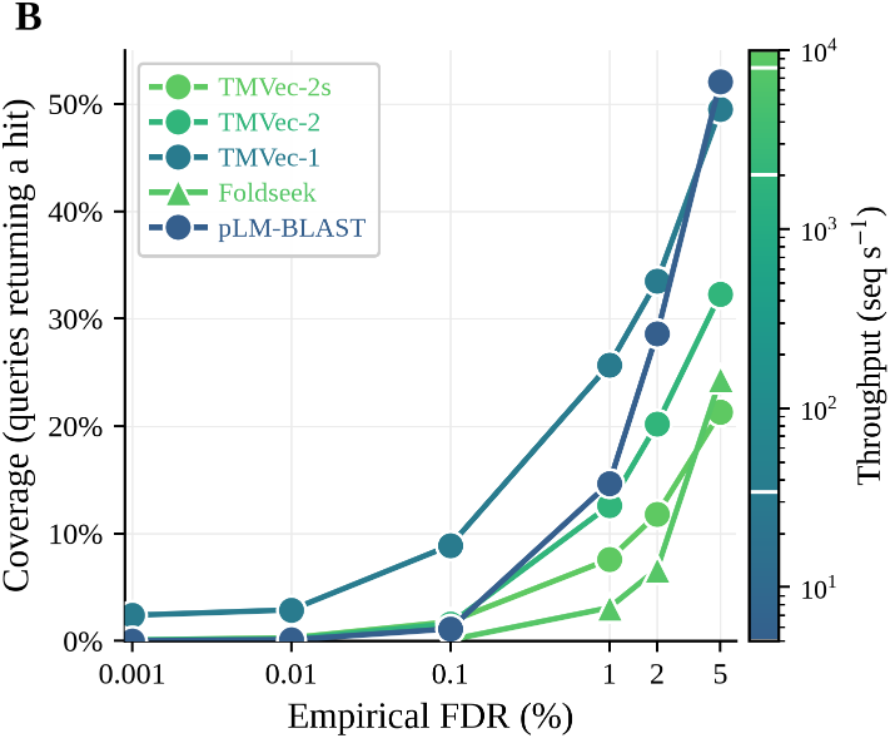
Retrieval coverage versus empirical false discovery rate for TM-Vec models. (target FDR *α ϵ* 0.001–0.2). TM-Vec 2 and 2s achieve performance close to the other benchmarked models at higher throughput.

At a 2% FDR, TM-Vec 2s achieves 12% coverage, Retrieving more hits than Foldseek and performing similarly to TM-Vec 2 and the original TM-Vec model. This indicates that conformal guaranties remain in place across various FDR thresholds, and showcases TM-Vec 2s provides high coverage rates at exponentially higher speeds than alternatives.

### Benchmark datasets

We evaluated all models on the CATH S100 [48] and SCOPe 40 datasets [49]. CATH S100 is a non-redundant subset of the CATH protein domain database. Specifically, we randomly sampled 10,000 protein domains from the CATH S100 dataset that were absent from training data. These represent five classes, 40 architectures, 769 topologies, and 1,765 superfamilies. We adopted the SCOPe 40 dataset from Foldseek’s benchmark [12], which in turn was derived from SCOPe 2.01 [49] by clustering at 40% sequence identity, resulting in 11,211 non-redundant protein structures. The dataset consists of manually classified single-domain proteins with an average length of 174 residues, organized into a hierarchy of seven classes, 1,194 folds, 1,960 superfamilies, and 4,161 families. Benchmarking was performed using a computer system with an AMD EPYC 7413 CPU and an NVIDIA A100 GPU (80GB VRAM).

## Code and data availability

The TM-Vec 2 and TM-Vec 2s models are available on Huggingface at https://huggingface.co/scikit-bio/tmvec-2 and https://huggingface.co/scikit-bio/tmvec-2s, respectively. The code and test datasets for bench-marking the models are available at: https://github.com/paarth-b/tmvec-bench.

## Author contributions

A.K.: Software, training and writing for TM-Vec 2s model. Helped P.B. in motivation experiments and benchmarking. P.B.: Software and training for TM-Vec 2 model. Conducted main experimentation for foundation model selection. V.B.: Dataset curation, software, experiment conceptualization, reviewing and editing. Q.Z. and J.M.: Conceptualization, methodology, resource acquisition, investigation, reviewing and editing.

## Acknowledgments

This research was supported by the U.S. Department of Energy, Office of Science under award DE-SC0024320. This work utilized computational resources of the DeltaAI supercomputer at the University of Illinois Urbana-Champaign, through the National Artificial Intelligence Research Resource (NAIRR) Pilot program (award: NAIRR250131), and computational resources of the Sol supercomputer at Arizona State University.

