## Supplementary materials for "TM-Vec 2s: Accelerated Protein Remote Homology Detection"

### Supplementary Figures

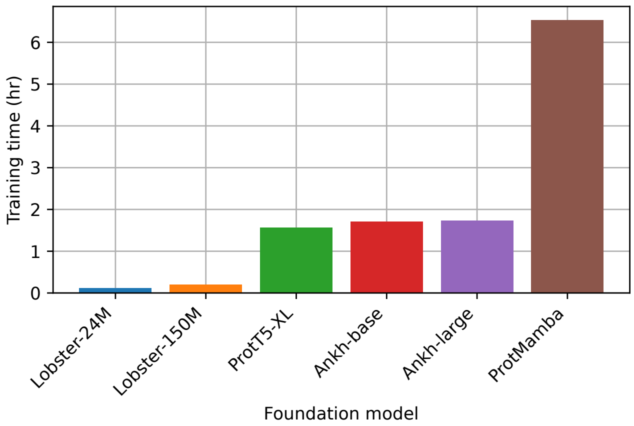


**Figure S1:** Fine-tuning times for state-of-the-art foundation protein language models on 10,000 sequences and two NVIDIA A100 GPUs.

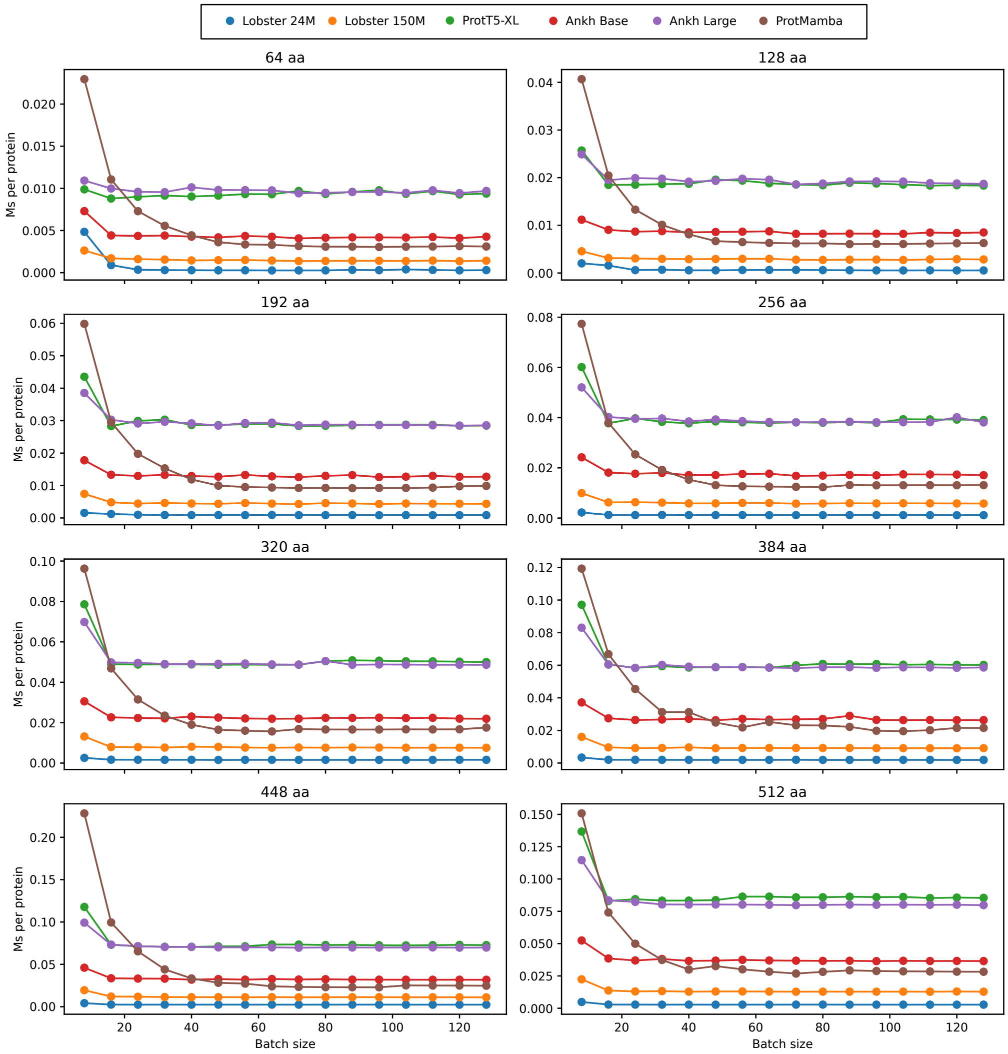


**Figure S2:** Inference times for state-of-the-art foundation protein language models with one NVIDIA A100 GPU. Eight distinct sequence length categories were tested: 64, 128, 192, 256, 320, 384, 448, and 512 amino acids. These lengths represent the typical range of protein sequences encountered in structural biology applications. Unit: milliseconds (Ms) per protein sequence.

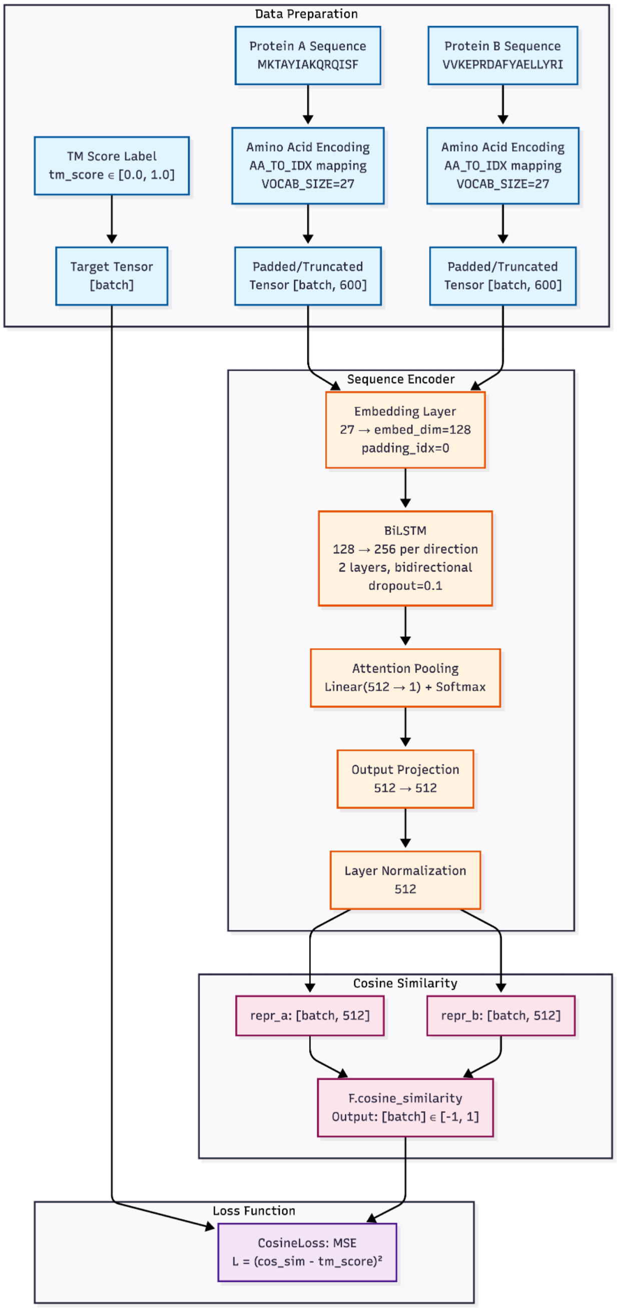


**Figure S3:** Student Model Architecture

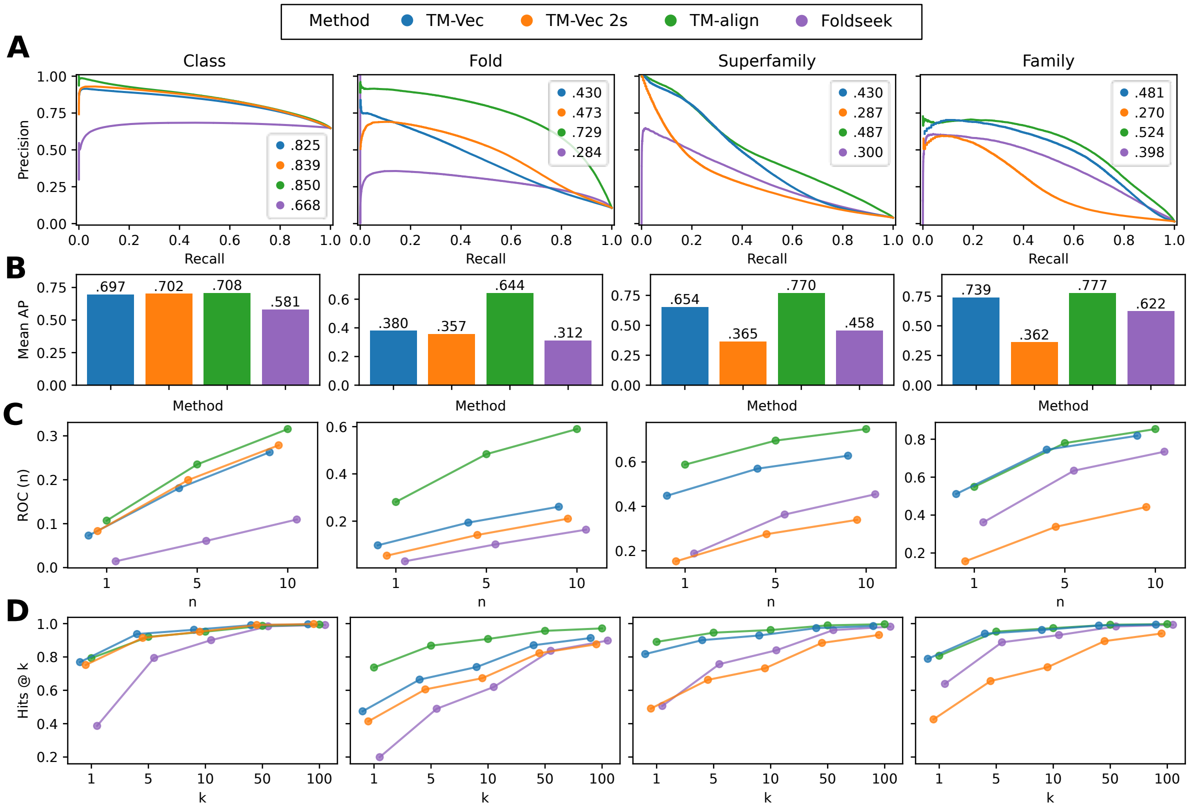


**Figure S4:** Comparison of homology detection performance of methods using the SCOPe 40 dataset with query-subject pairs filtered to those reported by all methods. **A**. PR curve and AP. **B**. Mean AP. **C**. $\mathrm{ROC_n}$. **D**. Hits@$k$. Refer to the caption of Fig. 3 for detailed description of the metrics.

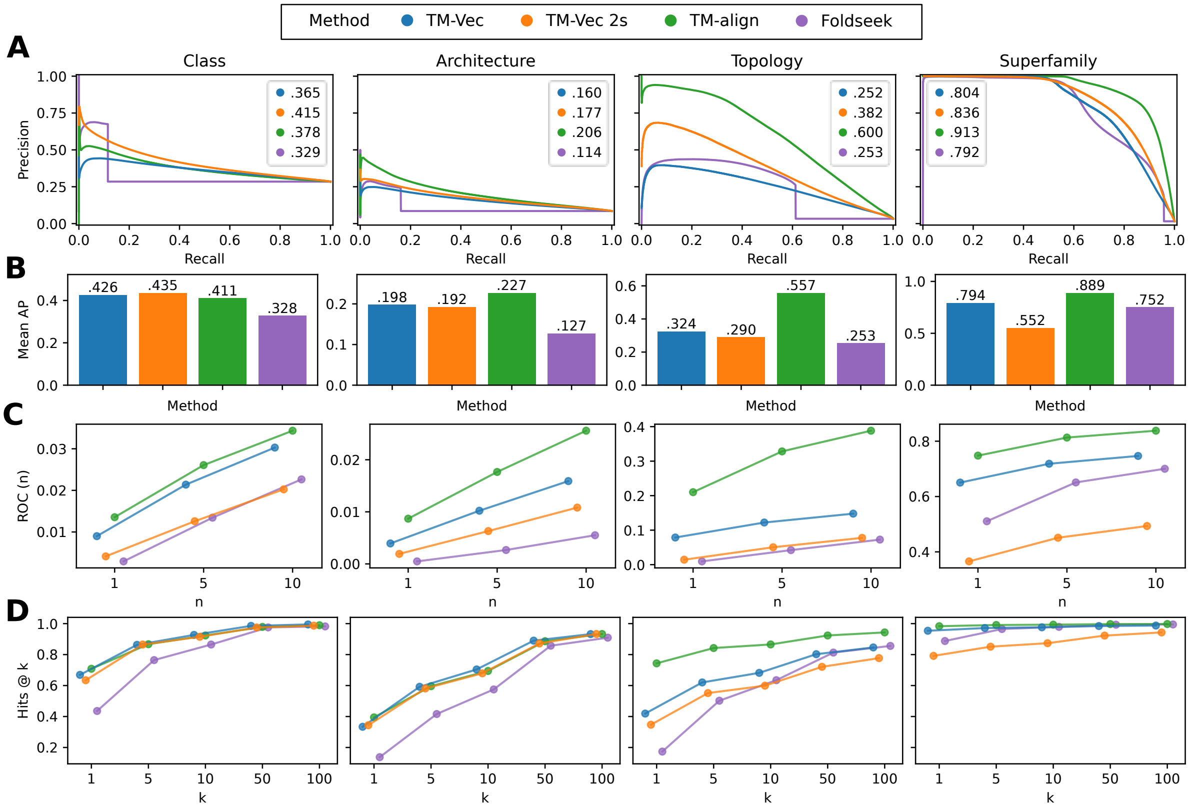


**Figure S5:** Comparison of homology detection performance of methods using the CATH 100 dataset with all query-subject pairs. **A**. PR curve and AP. **B**. Mean AP. **C**. $ROC_n$. **D**. Hits@$k$. Refer to the caption of Fig. 3 for detailed description of the metrics.

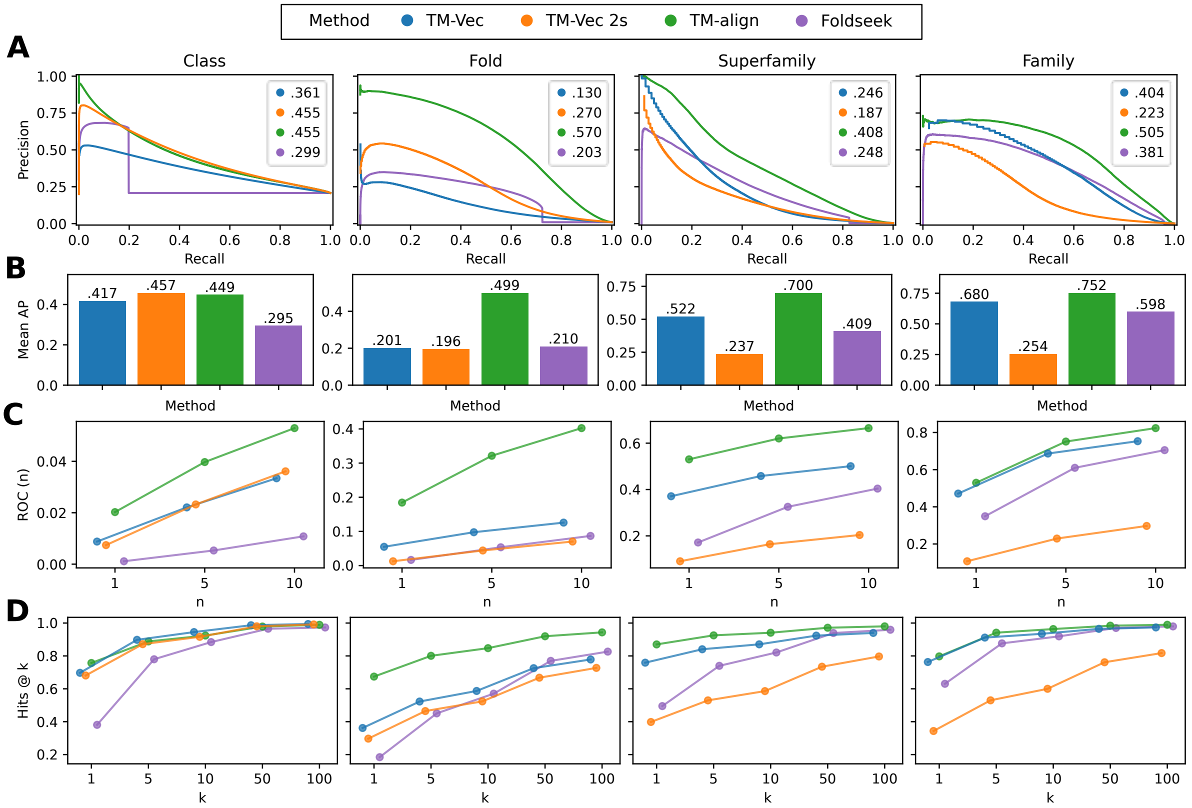


**Figure S6:** Comparison of homology detection performance of methods using the SCOPe 40 dataset with all query-subject pairs. **A**. PR curve and AP. **B**. Mean AP. **C**. $\mathrm{ROC_n}$. **D**. Hits@$k$. Refer to the caption of Fig. 3 for detailed description of the metrics.

### Supplementary Tables

**Table S1:** Encoding speed of TM-Vec models and existing methods across different sequence database sizes. Values represent runtime in seconds. Bold indicates best performance per setting.

| Sequences | TM-Vec | TM-Vec 2 | TM-Vec 2s | Foldseek | Foldseek GPU | Foldseek + ProstT5 |
|---|---|---|---|---|---|---|
| 10 | 0.355 | 0.021 | **0.020** | 0.293 | 0.308 | 3.665 |
| 100 | 3.668 | 0.207 | **0.024** | 0.314 | 0.414 | 6.673 |
| 1000 | 36.817 | 2.620 | **0.196** | 0.871 | 3.126 | 31.908 |
| 5000 | 183.446 | 17.303 | **0.985** | 2.175 | 14.384 | 145.600 |
| 10000 | 367.466 | 26.887 | **1.963** | 4.363 | 28.074 | 294.590 |
| 50000 | 1867.914 | 146.344 | **9.656** | 21.109 | 137.645 | 1468.185 |

**Table S2:** Search speed of TM-Vec models and existing methods across different query and database sizes. Values represent runtime in seconds. For TM-Vec and Foldseek variants, values are the total time of encoding query sequences and searching them against a pre-encoded database. For DIAMOND, which is a sequence alignment tool, values are search time only. Bold indicates best performance per setting.

| Query Size | Database Size | TM-Vec | TM-Vec 2 | TM-Vec 2s | Foldseek | Foldseek GPU | Foldseek + ProstT5 | DIAMOND |
|---|---|---|---|---|---|---|---|---|
| 10 | 1000 | 0.357 | **0.019** | 0.020 | 2.193 | 2.800 | 6.900 | 0.849 |
| 10 | 10000 | 0.362 | **0.019** | 0.020 | 2.310 | 3.955 | 6.846 | 0.996 |
| 10 | 100000 | 0.362 | **0.020** | 0.021 | 2.940 | 4.038 | 6.856 | 1.785 |
| 100 | 1000 | 3.640 | 0.208 | **0.024** | 2.582 | 7.130 | 11.394 | 0.840 |
| 100 | 10000 | 3.711 | 0.208 | **0.024** | 3.119 | 18.382 | 12.697 | 1.085 |
| 100 | 100000 | 3.693 | 0.209 | **0.025** | 4.438 | 25.759 | 13.265 | 2.143 |
| 1000 | 1000 | 36.835 | 2.530 | **0.210** | 7.539 | 89.635 | 53.542 | 0.904 |
| 1000 | 10000 | 37.034 | 2.517 | **0.227** | 12.943 | 208.745 | 62.742 | 1.130 |
| 1000 | 100000 | 37.042 | 2.517 | **0.200** | 29.121 | 445.525 | 69.444 | 2.568 |

**Table S3:** TM-score prediction accuracy metrics

| | | | | | | | **AP at threshold** | | | | |
|---|---|---|---|---|---|---|---|---|---|---|---|
| **Dataset** | **Model** | $r$ | $\rho$ | **MAE** | **RMSE** | **QWK** | 0.17 | 0.3 | 0.4 | 0.5 | 0.6 |
| CATH S100 | TM-Vec | 0.687 | 0.501 | 0.066 | 0.084 | 0.439 | 0.963 | 0.328 | 0.189 | 0.423 | 0.694 |
| | TM-Vec 2s | 0.744 | 0.563 | 0.047 | 0.061 | 0.577 | 0.965 | 0.370 | 0.317 | 0.571 | 0.725 |
| SCOPe 40 | TM-Vec | 0.513 | 0.470 | 0.076 | 0.095 | 0.290 | 0.933 | 0.273 | 0.086 | 0.107 | 0.204 |
| | TM-Vec 2s | 0.623 | 0.572 | 0.050 | 0.064 | 0.437 | 0.936 | 0.328 | 0.170 | 0.211 | 0.189 |

**Table S4:** Homology detection performance metrics on the CATH S100 dataset with query-subject pairs shared across methods.

| | | | ROC (*n*) | | | Hits @ *k* | | | | |
|---|---|---|---|---|---|---|---|---|---|---|
| **Method** | **AP** | **MAP** | 1 | 5 | 10 | 1 | 5 | 10 | 50 | 100 |
| **Class** (queries=9,715, pairs=4,234,614) | | | | | | | | | | |
| TM-Vec | 0.799 | 0.709 | 0.142 | 0.301 | 0.400 | 0.764 | 0.929 | 0.969 | 0.996 | 0.997 |
| TM-Vec 2s | 0.806 | 0.702 | 0.115 | 0.280 | 0.382 | 0.745 | 0.923 | 0.959 | 0.995 | 0.997 |
| TM-align | 0.780 | 0.684 | 0.155 | 0.294 | 0.380 | 0.758 | 0.915 | 0.954 | 0.991 | 0.996 |
| Foldseek | 0.672 | 0.602 | 0.030 | 0.129 | 0.219 | 0.441 | 0.776 | 0.879 | 0.992 | 0.998 |
| **Architecture** (queries=9,498, pairs=5,386,730) | | | | | | | | | | |
| TM-Vec | 0.366 | 0.369 | 0.031 | 0.093 | 0.148 | 0.386 | 0.682 | 0.787 | 0.916 | 0.947 |
| TM-Vec 2s | 0.322 | 0.344 | 0.024 | 0.078 | 0.126 | 0.403 | 0.679 | 0.774 | 0.931 | 0.964 |
| TM-align | 0.376 | 0.388 | 0.048 | 0.117 | 0.172 | 0.419 | 0.662 | 0.773 | 0.925 | 0.957 |
| Foldseek | 0.264 | 0.285 | 0.008 | 0.038 | 0.070 | 0.142 | 0.429 | 0.593 | 0.885 | 0.938 |
| **Topology** (queries=6,273, pairs=5,147,947) | | | | | | | | | | |
| TM-Vec | 0.620 | 0.532 | 0.130 | 0.232 | 0.303 | 0.534 | 0.773 | 0.831 | 0.928 | 0.954 |
| TM-Vec 2s | 0.647 | 0.479 | 0.055 | 0.158 | 0.237 | 0.477 | 0.696 | 0.760 | 0.878 | 0.923 |
| TM-align | 0.800 | 0.712 | 0.301 | 0.481 | 0.570 | 0.801 | 0.896 | 0.921 | 0.972 | 0.984 |
| Foldseek | 0.402 | 0.388 | 0.018 | 0.079 | 0.134 | 0.191 | 0.553 | 0.699 | 0.895 | 0.942 |
| **Superfamily** (queries=9,095, pairs=7,859,443) | | | | | | | | | | |
| TM-Vec | 0.896 | 0.867 | 0.704 | 0.785 | 0.819 | 0.969 | 0.984 | 0.988 | 0.996 | 0.998 |
| TM-Vec 2s | 0.893 | 0.666 | 0.444 | 0.559 | 0.614 | 0.843 | 0.908 | 0.931 | 0.971 | 0.983 |
| TM-align | 0.941 | 0.920 | 0.780 | 0.850 | 0.875 | 0.987 | 0.995 | 0.996 | 0.999 | 0.999 |
| Foldseek | 0.826 | 0.784 | 0.527 | 0.676 | 0.730 | 0.891 | 0.969 | 0.983 | 0.997 | 0.998 |

**Table S5:** Homology detection performance metrics on the SCOPe 40 dataset with query-subject pairs shared across methods.

| | | | **ROC (*n*)** | | | **Hits @ *k*** | | | | |
|---|---|---|---|---|---|---|---|---|---|---|
| **Method** | **AP** | **MAP** | 1 | 5 | 10 | 1 | 5 | 10 | 50 | 100 |
| **Class** (queries=10,976, pairs=7,909,702) | | | | | | | | | | |
| TM-Vec | 0.825 | 0.697 | 0.073 | 0.181 | 0.263 | 0.769 | 0.937 | 0.963 | 0.991 | 0.996 |
| TM-Vec 2s | 0.839 | 0.702 | 0.083 | 0.199 | 0.279 | 0.752 | 0.914 | 0.951 | 0.992 | 0.997 |
| TM-align | 0.850 | 0.708 | 0.107 | 0.235 | 0.316 | 0.793 | 0.921 | 0.952 | 0.987 | 0.994 |
| Foldseek | 0.668 | 0.581 | 0.014 | 0.061 | 0.109 | 0.387 | 0.794 | 0.901 | 0.984 | 0.992 |
| **Fold** (queries=5,040, pairs=3,958,982) | | | | | | | | | | |
| TM-Vec | 0.430 | 0.380 | 0.098 | 0.194 | 0.261 | 0.474 | 0.664 | 0.740 | 0.870 | 0.913 |
| TM-Vec 2s | 0.473 | 0.357 | 0.054 | 0.142 | 0.210 | 0.414 | 0.606 | 0.673 | 0.823 | 0.876 |
| TM-align | 0.729 | 0.644 | 0.281 | 0.484 | 0.590 | 0.737 | 0.868 | 0.908 | 0.957 | 0.971 |
| Foldseek | 0.284 | 0.312 | 0.030 | 0.102 | 0.164 | 0.199 | 0.489 | 0.620 | 0.837 | 0.898 |
| **Superfamily** (queries=8,438, pairs=7,220,100) | | | | | | | | | | |
| TM-Vec | 0.430 | 0.654 | 0.448 | 0.570 | 0.628 | 0.817 | 0.901 | 0.929 | 0.973 | 0.985 |
| TM-Vec 2s | 0.287 | 0.365 | 0.152 | 0.275 | 0.339 | 0.491 | 0.663 | 0.732 | 0.884 | 0.932 |
| TM-align | 0.487 | 0.770 | 0.588 | 0.696 | 0.748 | 0.890 | 0.946 | 0.961 | 0.990 | 0.996 |
| Foldseek | 0.300 | 0.458 | 0.188 | 0.363 | 0.455 | 0.506 | 0.757 | 0.840 | 0.961 | 0.981 |
| **Family** (queries=8,849, pairs=7,230,346) | | | | | | | | | | |
| TM-Vec | 0.481 | 0.739 | 0.511 | 0.745 | 0.818 | 0.789 | 0.940 | 0.962 | 0.988 | 0.994 |
| TM-Vec 2s | 0.270 | 0.362 | 0.156 | 0.338 | 0.442 | 0.426 | 0.655 | 0.739 | 0.895 | 0.941 |
| TM-align | 0.524 | 0.777 | 0.549 | 0.780 | 0.854 | 0.807 | 0.952 | 0.973 | 0.993 | 0.997 |
| Foldseek | 0.398 | 0.622 | 0.362 | 0.634 | 0.734 | 0.638 | 0.888 | 0.931 | 0.983 | 0.993 |

**Table S6:** Homology detection performance metrics on the CATH S100 dataset with all query-subject pairs.

| | | | **ROC (*n*)** | | | **Hits @ *k*** | | | | |
|---|---|---|---|---|---|---|---|---|---|---|
| **Method** | **AP** | **MAP** | 1 | 5 | 10 | 1 | 5 | 10 | 50 | 100 |
| **Class** (queries=9,912, pairs=87,429,562) | | | | | | | | | | |
| TM-Vec | 0.365 | 0.426 | 0.009 | 0.021 | 0.030 | 0.669 | 0.863 | 0.927 | 0.985 | 0.995 |
| TM-Vec 2s | 0.415 | 0.435 | 0.004 | 0.013 | 0.020 | 0.634 | 0.865 | 0.915 | 0.975 | 0.987 |
| TM-align | 0.378 | 0.411 | 0.014 | 0.026 | 0.034 | 0.708 | 0.867 | 0.923 | 0.979 | 0.990 |
| Foldseek | 0.329 | 0.328 | 0.003 | 0.013 | 0.023 | 0.436 | 0.764 | 0.865 | 0.976 | 0.982 |
| **Architecture** (queries=9,803, pairs=94,361,397) | | | | | | | | | | |
| TM-Vec | 0.160 | 0.198 | 0.004 | 0.010 | 0.016 | 0.334 | 0.592 | 0.704 | 0.890 | 0.932 |
| TM-Vec 2s | 0.177 | 0.192 | 0.002 | 0.006 | 0.011 | 0.344 | 0.581 | 0.679 | 0.872 | 0.933 |
| TM-align | 0.206 | 0.227 | 0.009 | 0.018 | 0.026 | 0.394 | 0.596 | 0.694 | 0.886 | 0.933 |
| Foldseek | 0.114 | 0.127 | 0.000 | 0.003 | 0.006 | 0.138 | 0.416 | 0.575 | 0.857 | 0.909 |
| **Topology** (queries=6,910, pairs=67,771,198) | | | | | | | | | | |
| TM-Vec | 0.252 | 0.324 | 0.079 | 0.122 | 0.148 | 0.419 | 0.620 | 0.682 | 0.802 | 0.846 |
| TM-Vec 2s | 0.382 | 0.290 | 0.014 | 0.050 | 0.078 | 0.347 | 0.551 | 0.599 | 0.721 | 0.777 |
| TM-align | 0.600 | 0.557 | 0.210 | 0.328 | 0.388 | 0.743 | 0.842 | 0.865 | 0.923 | 0.943 |
| Foldseek | 0.253 | 0.253 | 0.009 | 0.042 | 0.073 | 0.174 | 0.502 | 0.635 | 0.813 | 0.856 |
| **Superfamily** (queries=9,153, pairs=91,520,847) | | | | | | | | | | |
| TM-Vec | 0.804 | 0.794 | 0.650 | 0.718 | 0.747 | 0.954 | 0.971 | 0.976 | 0.984 | 0.987 |
| TM-Vec 2s | 0.836 | 0.552 | 0.366 | 0.451 | 0.493 | 0.792 | 0.851 | 0.873 | 0.923 | 0.943 |
| TM-align | 0.913 | 0.889 | 0.748 | 0.813 | 0.838 | 0.983 | 0.991 | 0.993 | 0.997 | 0.997 |
| Foldseek | 0.792 | 0.752 | 0.511 | 0.651 | 0.700 | 0.888 | 0.965 | 0.979 | 0.993 | 0.995 |

**Table S7:** Homology detection performance metrics on the SCOPe 40 dataset with all query-subject pairs.

| | | | **ROC (*n*)** | | | **Hits @ *k*** | | | | |
|---|---|---|---|---|---|---|---|---|---|---|
| **Method** | **AP** | **MAP** | 1 | 5 | 10 | 1 | 5 | 10 | 50 | 100 |
| **Class** (queries=11,211, pairs=124,638,772) | | | | | | | | | | |
| TM-Vec | 0.361 | 0.417 | 0.009 | 0.022 | 0.033 | 0.697 | 0.898 | 0.945 | 0.987 | 0.994 |
| TM-Vec 2s | 0.455 | 0.457 | 0.007 | 0.023 | 0.036 | 0.681 | 0.872 | 0.916 | 0.980 | 0.992 |
| TM-align | 0.455 | 0.449 | 0.020 | 0.040 | 0.053 | 0.757 | 0.888 | 0.924 | 0.979 | 0.989 |
| Foldseek | 0.299 | 0.295 | 0.001 | 0.005 | 0.011 | 0.381 | 0.780 | 0.884 | 0.965 | 0.973 |
| **Fold** (queries=5,488, pairs=61,302,578) | | | | | | | | | | |
| TM-Vec | 0.130 | 0.201 | 0.055 | 0.098 | 0.126 | 0.362 | 0.523 | 0.587 | 0.726 | 0.778 |
| TM-Vec 2s | 0.270 | 0.196 | 0.013 | 0.044 | 0.070 | 0.297 | 0.465 | 0.524 | 0.667 | 0.727 |
| TM-align | 0.570 | 0.499 | 0.184 | 0.321 | 0.402 | 0.674 | 0.801 | 0.847 | 0.920 | 0.943 |
| Foldseek | 0.203 | 0.210 | 0.017 | 0.054 | 0.087 | 0.184 | 0.451 | 0.571 | 0.770 | 0.826 |
| **Superfamily** (queries=8,636, pairs=96,717,780) | | | | | | | | | | |
| TM-Vec | 0.246 | 0.522 | 0.371 | 0.458 | 0.501 | 0.759 | 0.841 | 0.871 | 0.923 | 0.940 |
| TM-Vec 2s | 0.187 | 0.237 | 0.091 | 0.164 | 0.204 | 0.398 | 0.530 | 0.586 | 0.734 | 0.797 |
| TM-align | 0.408 | 0.700 | 0.530 | 0.620 | 0.665 | 0.870 | 0.925 | 0.941 | 0.971 | 0.980 |
| Foldseek | 0.248 | 0.409 | 0.172 | 0.325 | 0.404 | 0.495 | 0.740 | 0.821 | 0.939 | 0.959 |
| **Family** (queries=8,975, pairs=100,609,750) | | | | | | | | | | |
| TM-Vec | 0.404 | 0.680 | 0.471 | 0.687 | 0.753 | 0.763 | 0.912 | 0.935 | 0.965 | 0.973 |
| TM-Vec 2s | 0.223 | 0.254 | 0.107 | 0.229 | 0.297 | 0.343 | 0.530 | 0.600 | 0.762 | 0.817 |
| TM-align | 0.505 | 0.752 | 0.529 | 0.751 | 0.824 | 0.797 | 0.941 | 0.963 | 0.984 | 0.989 |
| Foldseek | 0.381 | 0.598 | 0.349 | 0.610 | 0.705 | 0.630 | 0.876 | 0.919 | 0.970 | 0.980 |
